# Feature-based attention induces surround suppression during the perception of visual motion

**DOI:** 10.1101/2021.02.17.431646

**Authors:** Sang-Ah Yoo, Julio C. Martinez-Trujillo, Stefan Treue, John K. Tsotsos, Mazyar Fallah

## Abstract

Attention to a stimulus feature prioritizes its processing while strongly suppressing the processing of similar features, a non-linear phenomenon called surround suppression. Here we investigated this phenomenon using neurophysiology and psychophysics. We recorded responses of motion direction-selective neurons in area MT/MST of monkeys in different conditions. When attention was allocated to a stimulus moving in the neurons’ preferred direction responses to a distractor were strongly suppressed for directions nearby the preferred direction. These effects were modeled as the interaction between two Gaussian fields representing narrowly-tuned excitatory and widely-tuned inhibitory inputs into a neuron, with attention more strongly modulating the gain of the inhibitory inputs. We additionally demonstrated a corresponding behavioral effect in humans: Feature-based attention strongly reduced motion repulsion in the vicinity of the attended motion direction. Our results demonstrate that feature-based attention can induce non-linear changes in neuronal tuning curves via unbalanced gain changes to excitatory and inhibitory inputs into neurons ultimately translating into similar effects during behavior.

## Introduction

Attention, defined as the selection and modulation of information processing in the brain, allows sensory systems to deal with information processing overload^1^. Attention can be allocated to a region of space (spatial attention) or to the features of an object (feature-based attention). Feature-based attention (FBA) facilitates and prioritizes the processing of the attended feature relative to unattended ones, as corroborated by a number of studies using various methodologies ^2–11^. Electrophysiological studies in behaving macaque monkeys have shown that FBA can increase or decrease the responses of sensory neurons to visual stimuli ^4,12^. This effect was described as a monotonic change in the gain of neuronal responses following the feature-similarity gain principle ^4^. However, at least one computational model (the Selective Tuning model (ST) ^1,13,14^) has predicted that the suppressive effect of FBA is greater for features more similar to the attended feature relative to those that are dissimilar. Such non-linear feature-based surround suppression has been also observed in human studies ^15–22^. The neurophysiological substrates of feature-based surround suppression remain elusive.

Electrophysiological studies in macaques reported that FBA maximally enhances responses of neurons selective for the attended feature, and that such enhancement grows smaller to become a suppression as the neuron’s preferred feature differs from the attended feature ^4,8^. On the other hand, ST proposes that when a stimulus is attended responses to nearby unattended stimuli are suppressed. Since in real scenes most attended stimuli have other stimuli nearby (the *context problem* ^14^), ST ameliorates contextual interference via top-down attention, e.g., suppressing responses to stimuli in the neighborhood of the attended stimulus. The context problem and ST’s solution can apply to space, features, or objects ^1,13,14^. However, this proposal has not been supported by subsequent neurophysiological studies of FBA that instead, document a monotonic modulation ^4,8^.

Behavioral studies used a visual motion attentional cueing paradigm found that participants’ performance decreased as the direction offset between a cue and a target stimulus became greater; however, performance gradually recovered when the offset was larger than 90 deg ^20,21^, indicating feature-based surround suppression in motion processing. On the other hand, a previous neurophysiological study of FBA has shown that tuning curves of direction-selective neurons in the middle temporal area (MT) show changes in gain, baseline, and width, but tuning curves keep their monotonic Gaussian profiles ^4^. This study, however, used moving random dot patterns (RDPs) positioned in different hemifields and recorded from area MT neurons with receptive fields (RFs) localized to the contralateral visual hemifield. It is possible that the lack of interference (context within the ST framework) due to the stimuli being far away and the relatively small attentional modulation of responses documented in these conditions is insufficient to isolate the feature-based surround suppression predicted by ST and observed in behavioral studies.

The present study aims to investigate feature-based surround suppression at neurophysiological and behavioral levels. First, we measured the activity of direction-selective neurons in MT and medial superior temporal (MST) visual cortical areas of macaque monkeys. We obtained tuning curves of direction-selective neurons by placing two moving random dot patterns (RDPs) within a neuron’s RF – one RDP always moved in the neuron’s preferred direction (preferred pattern) and the other moved in one of twelve different directions (tuning pattern). In different trials, we instructed the animals to direct attention either to the fixation point (fixation condition) or to one of the RDPs (attend-preferred and attend-tuning conditions). We found that during fixation neuronal and population tuning curves were well fitted by a single Gaussian curve with positive gain. However, when the animals attended to the preferred pattern neuronal tuning curves exhibited a suppressive surround. Here, response profiles were better described by adding a second wider Gaussian function with negative gain. We modeled the feature-based surround suppression by the additive interaction of the two Gaussian fields representing excitatory and inhibitory input fields into a neuron. FBA disproportionally increases the gain of the inhibitory wider relative to the excitatory narrower input field producing a suppressive surround.

In the behavioral experiment, we measured behavioral correlates of this FBA suppressive effect on motion repulsion. Motion repulsion is a perceptual illusion arising from an overestimation of the directional difference between the two superimposed motion surfaces ^23–25^. FBA modulates motion repulsion ^26,27^. We measured motion repulsion under different attentional conditions (focused vs. divided attention) while the directional difference between two motion surfaces varied. We observed FBA caused surround suppression on motion repulsion.

## Results

### Neurophysiology

Two macaque monkeys were trained to selectively attend to a cued stimulus, while keeping gaze on a fixation point (Figure 1A). We positioned two RDPs within a neuron’s RF, one RDP always moved in the neuron’s preferred direction (preferred pattern) and the other could move in one of 12 different directions from trial to trial (tuning pattern, in steps of 30 deg). The animals attended to and reported a direction change either in the preferred pattern (attend-preferred condition) or tuning pattern (attend-tuning condition) (Figure 1B). In the fixation condition they detected a color change of the fixation point while ignoring both RDPs. Animal F and M achieved 86% and 87% of change detection accuracy, respectively, indicating that they correctly perform the task.

**Figure 1.**
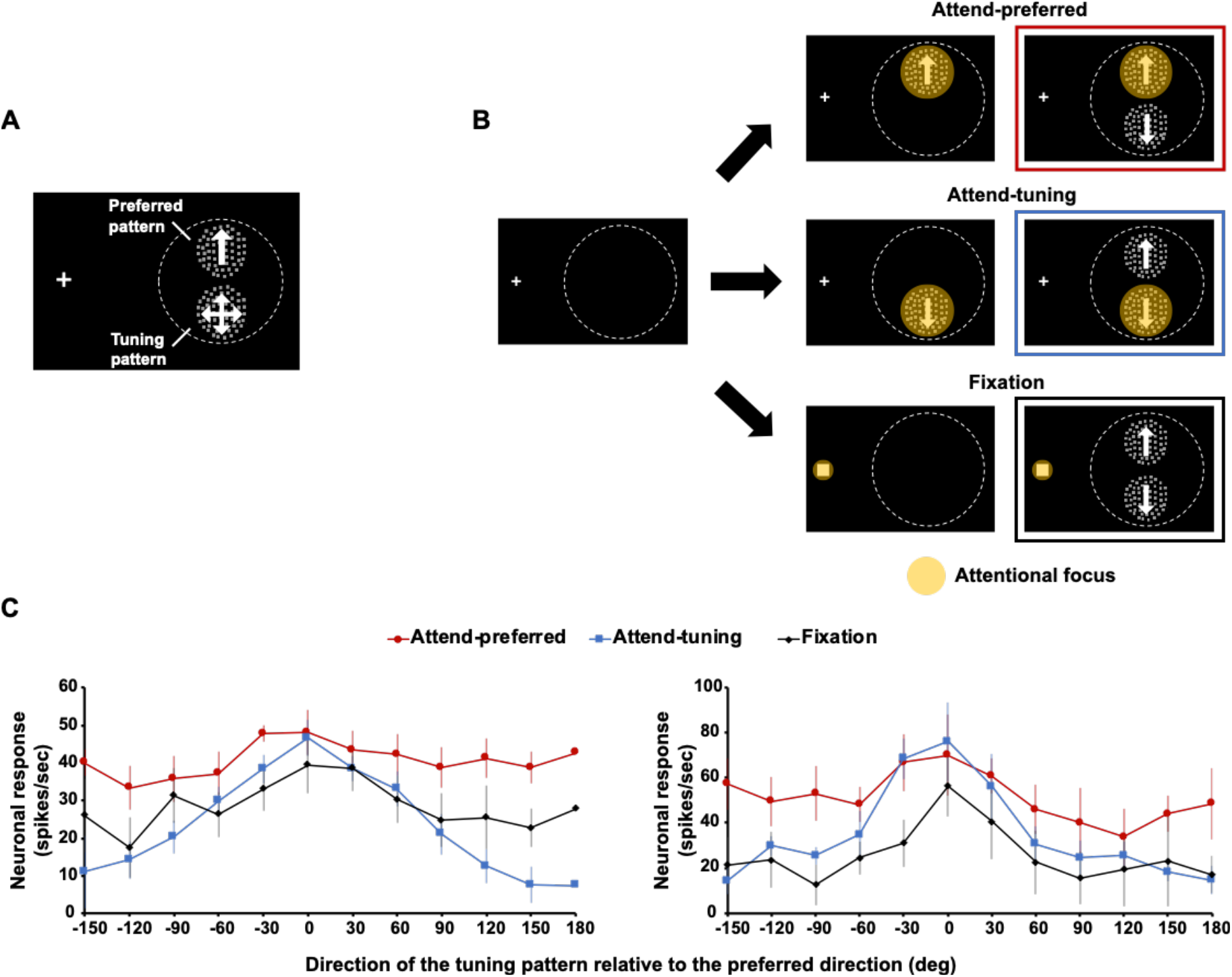
(A) Stimulus configuration of Experiment 1. The preferred pattern (always moving in the neuron’s preferred direction) and the tuning pattern (moving in one of 12 directions) were presented within a neuron’s RF. The fixation point was presented on the left side of the display. (B) Three attentional conditions. While an animal foveated the fixation point, it was cued to attend to either the preferred (attend-preferred) or tuning (attend-tuning) pattern and reported a directional change of the attended pattern. In the fixation condition the animal was cued to the fixation point and had to report a change in the color of a small square superimposed on the fixation point while ignoring the RDPs. The yellow spot indicates the allocation of attention in each condition. (C) Responses of two single neurons in different attentional conditions. The abscissa represents the direction of the tuning pattern as a function of the distance to the preferred direction and the ordinate represents the response in spikes/second. Error bars indicate the standard deviation (SD).

Seventy-eight neurons were included in the analysis. Two examples of neuronal responses in different attentional conditions are shown in Figure 1C. Responses in the attend-preferred condition were in general stronger than in the other conditions (labelled as zero in the abscissa of Figure 1C). In order to determine the difference in responses between conditions we normalized responses in each neuron to the maximal response, we then align all the responses to the direction producing the maximal response and pooled across neurons for each condition. We aligned the responses along the x-axis as a function of the difference between the preferred direction and the direction of the tuning pattern in a symmetrical manner to produce a response profile (Figure 2A).

**Figure 2.**
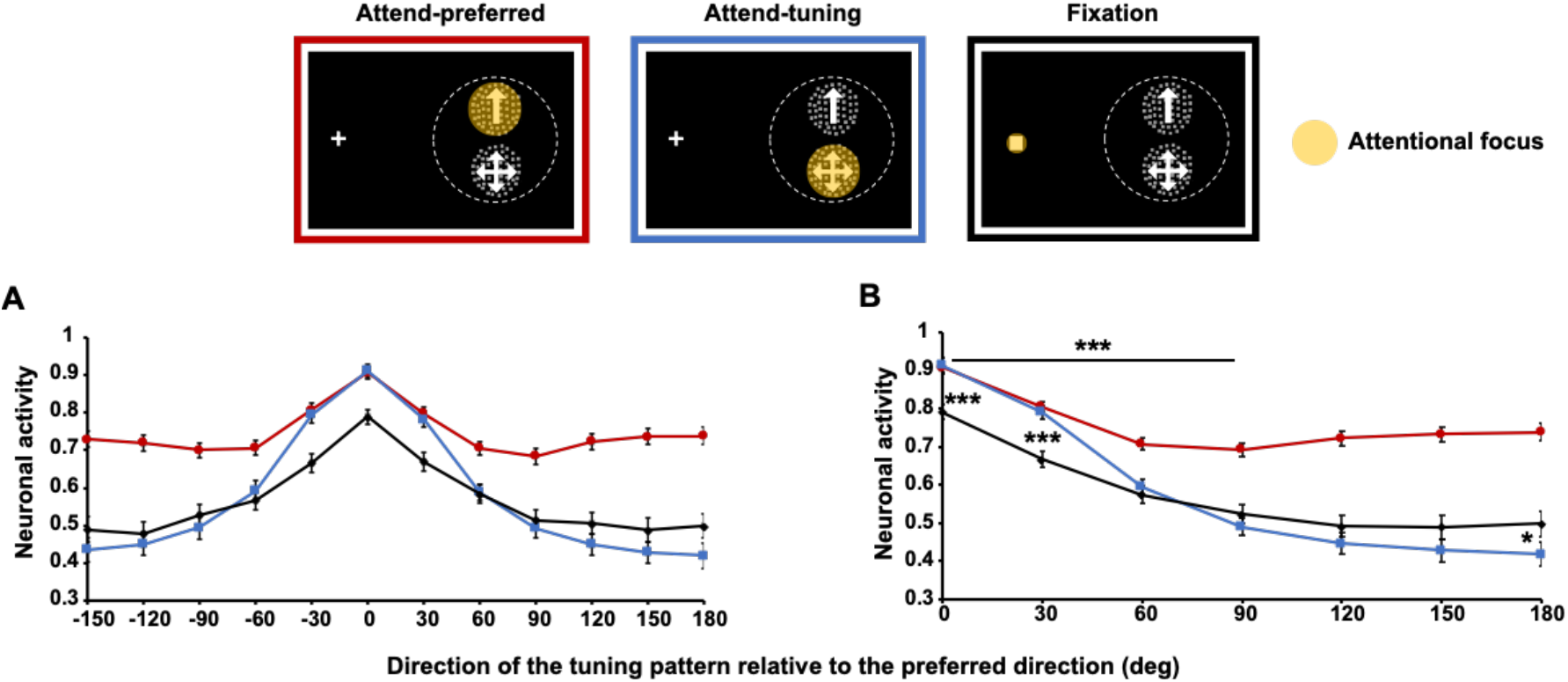
(A) Average normalized neuronal response in each attentional condition. (B) Normalized neuronal responses averaged across the same (absolute) directional difference. Responses in the attend-tuning and the fixation conditions monotonically decrease as the offset between the preferred and tuning patterns increases. Responses in the attend-tuning condition are larger than that in the fixation condition when the direction of the tuning pattern is close to the neuron’s preferred direction (0~30 deg). This relationship reverses when the direction of the tuning pattern approaches the neuron’s anti-preferred direction. In the attend-preferred condition, however, responses are lowest when the direction of the tuning pattern is approximately 90 deg away from the preferred direction. Then, the responses tend to increase as the directional difference increases (~180 deg difference, *p* = .051). Error bars indicate standard error of the mean (SEM) across the normalized responses of each neuron. * *p* < .05, ** *p* < .01, *** *p* < .001

A repeated-measures ANOVA showed that the neuronal response was significantly modulated by attentional conditions (*F*(2, 122) = 47.058, *p* < .001). There was an increased overall response in the attend-preferred condition relative to the other conditions (all *p*s < .001, Bonferroni correction was used for all multiple comparisons). No significant difference was observed between the attend-tuning and the fixation conditions (*p* = 1). The direction of the tuning pattern relative to the preferred direction also significantly modulated neuronal responses (*F*(11, 671) = 45.768, *p* < .001), with the greatest response when the tuning pattern moved in the preferred direction (0 deg difference, all *p*s < .001) and a gradual decrease as the tuning pattern’s direction deviated from the preferred direction. The interaction between attentional conditions and directional difference was significant (*F*(22, 1342) = 13.561, *p* < .001). To elucidate the nature of this interaction, we examined the profile of attentional modulation under different attentional conditions. Because the response functions show a symmetric profile, we collapsed neuronal responses when the absolute directional difference between the preferred and the tuning patterns was the same (e.g., ±30 deg) (Figure 2B).

Neuronal responses in the fixation condition peaked when the tuning pattern moved in the neuron’s preferred direction likely because both RDPs moved in the preferred direction and neither pattern extended into the RF’s inhibitory surround. This was assessed during initial mapping of the RF ^4^. The response reached its minimum when the tuning pattern moved in the neuron’s anti-preferred direction. There was a monotonic decrease in response as a function of the difference between the neuron’s preferred direction and the direction of the tuning pattern (Figure 2B).

Neuronal response in the attend-tuning condition also monotonically decreased as the directions between the preferred and the tuning patterns became dissimilar. In this condition, neuronal responses were greater than those in the fixation condition when the direction of the tuning pattern was closer to the neuron’s preferred direction (at 0 and 30 deg differences, all *p*s < .001). On the other hand, responses in the attend-tuning condition were lower than those in the fixation condition when the tuning pattern moved in the neuron’s anti-preferred direction (M_diff_ = −.07, SE = .04, *t*(76) = −2.101, *p* = .039). This effect is similar to feature-similarity gain modulation described in other studies ^4^.

In the attend-preferred condition, the maximal neuronal response was also observed when the directions of the preferred and tuning patterns were the same. However, we did not observe a monotonic response decrease with the direction of the tuning pattern relative to the preferred direction. Responses were lowest when the tuning pattern moved in directions ~90 deg away from the preferred direction, not when the tuning pattern moved in the anti-preferred direction. The minimum neuronal response was significantly lower than the maximum response (90 vs. 0 deg, M_diff_ = −.22, SE = .02, *t*(76) = −8.78, *p* < .001) and it was lower than the response for the anti-preferred direction (90 vs. 180 deg, M_diff_ = −.04, SE = .02, *t*(76) = −2.272, *p* = .051). This indicates that the attend-preferred condition causes a suppressive surround for motion quasi-orthogonal to a neuron’s preferred direction.

### Tuning curves

Neuron’s tuning curves have been modeled using Gaussian functions ^8^. However, because responses do not clearly follow a monotonic profile in the attend-preferred condition, we fitted two different models to the average normalized neuronal response (population response) in each experimental condition, a single Gaussian and a sum of two Gaussians. In the attend-preferred condition (Figure 3A), the sum of two Gaussians (adjusted R^2^ = .990) model explained the population response better than the single Gaussian model (adjusted R^2^ = .886). This was mainly due to the ability of the sum of two Gaussians (one Gaussian having a positive gain and narrower width than the second Gaussian with negative gain and larger width) to account for the decrease in response in the vicinity of the preferred direction relative to directions farther away from the preferred one. This effect resembles the feature-based surround suppression reported in previous behavioral and modeling studies ^15–22^. In the other attentional conditions (Figure 3B and 3C), both models could explain the response profiles equally well (all adjusted R^2^, attend-tuning condition: single Gaussian (.994), sum of two Gaussians (.999); fixation condition: single Gaussian (.957), sum of two Gaussians (.987)).

**Figure 3.**
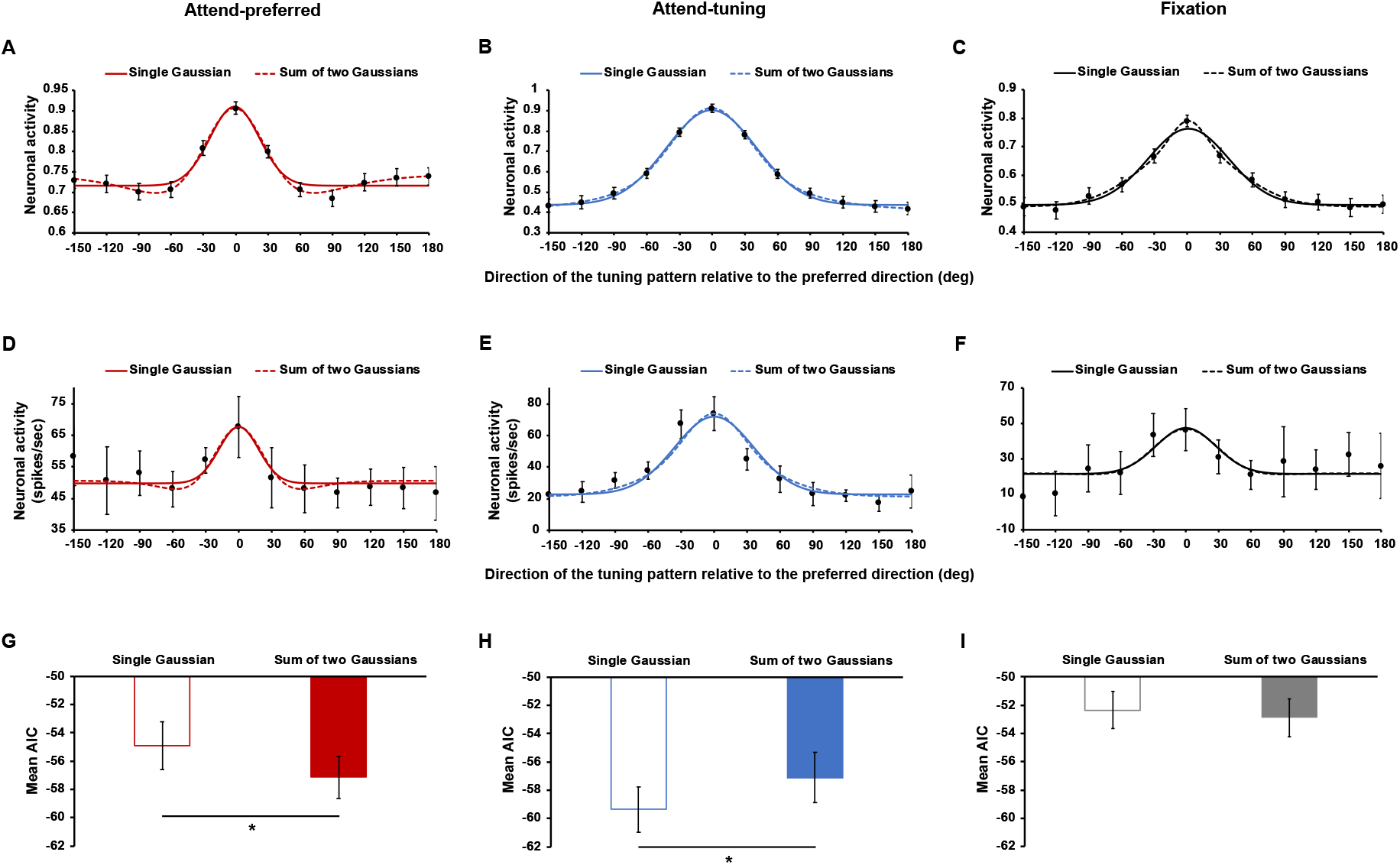
(A-C) Different model fits to the average normalized neuronal response (population level) in the attend-preferred, attend-tuning, and fixation conditions. Solid curve indicates the single Gaussian model fit and dashed curve indicates the sum of two Gaussians model fit (see D-F for an example neuron). (G-I) Akaike Information Criterion (AIC) for each model fit was measured at individual neuron level. Smaller AIC value means better model fit. Outlined and filled bars indicate the mean AIC of the single Gaussian and the sum of two Gaussians models, respectively. Error bars in Fig 3A-C, and 3G-I indicate SEM, and those in Fig 3D-F indicate SD. * *p* < .05

In order to quantify these observations, we compared model fits at the individual neuron level. We included individual neurons in the analysis only if both models reasonably fit their responses. We then compared different model fits to the same neuronal response (i.e., pairwise comparison): 69 neurons in the attend-preferred condition, 53 neurons in the attend-tuning condition, and 59 neurons in the fixation condition were included in the analysis. Example model fits to normalized responses are shown in Figure 3D-F. As a goodness-of-fit measure, we computed the Akaike Information Criteria (AIC) for each model fit. The AIC takes into account the number of parameters in each model, which is lower in the single Gaussian compared to the sum of two Gaussians. In the attend-preferred condition (Figure 3G), the AIC was greater for the single Gaussian than for the sum of two Gaussians (M_diff_ = 2.22, SE = 1.03, *t*(68) = 2.148, *p* = .035), meaning that the latter model explained the data better. Conversely, the single Gaussian fitted the data better in the attend-tuning condition (Figure 3H), demonstrating a smaller AIC value than the sum of two Gaussians (M_diff_ = −2.25, SE = .92, *t*(52) = −2.448, *p* = .018). In the fixation condition (Figure 3I), there was no significant difference in AIC between the two models (M_diff_ = .53, SE = .88, *t*(58) = .605, *p* = .548).

Overall, for the population as well as for single neurons the sum of two Gaussians model fits the data better only in the attend-preferred condition. In the other conditions, either both models fit the data equally well (fixation) or the single Gaussian model performs better than the sum of two Gaussians model (attend tuning). These results indicate that feature-based surround suppression became evident only in the attend-preferred condition.

### Modeling feature-based surround suppression

The attend-preferred condition may have allowed us to isolate the feature-based surround suppression effect because attention was always on the same feature (preferred direction) while the distractor tuning pattern changed direction from trial to trial. Thus, keeping attention on the preferred direction reveals the surround suppressive effect on distracting features. One possible explanation for this effect is the interaction between excitatory- and inhibitory-tuned inputs into a neuron during the allocation of FBA. In order to model this interaction, we modeled the excitatory and inhibitory fields of direction-selective neurons as two Gaussian functions with positive and negative gain, respectively. We first fit the average normalized neuronal responses in the fixation condition where no attention was directed to motion direction with the single Gaussian model:

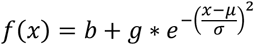

where *b* is the baseline, *g* is the gain of the response, μ is the position of the center of the maximum response, and σ is the width of the Gaussian model. μ is always 0 because neuronal response was maximized when the preferred and tuning pattern moved in the same direction (no directional difference).

An important point here is that we assumed that in the fixation condition, the predominant contribution to the response is provided by excitatory-tuned inputs into the cell with σ representing the width or selectivity of the inputs. The contribution of inhibitory inputs into the cell is considered small here and will be captured by the single Gaussian ^28^. The fitting result is illustrated in Figure 4A (yellow line). The coefficients and goodness-of-fit measures are shown in Table 1.

**Table 1.** Coefficients of the model fits to the fixation and attend-preferred conditions and goodness-of-fit measures.

|  |  | Fixation | Attend-preferred (constrained) | Attend-preferred (unconstrained) |
| --- | --- | --- | --- | --- |
| Baseline |  | 0.4962 | 0.7566 | 0.7438 |
| 1<sup>st</sup> Gaussian (excitatory) | Gain | 0.2713 | 0.2713 | 0.2774 |
|  | Width | 51.29 | 51.29 | 42.65 |
| 2<sup>nd</sup> Gaussian (inhibitory) | Gain | N/A | -0.1468 | -0.1168 |
|  | Width |  | 110.8 | 96.94 |
| Goodness-of-fit | SSE | 0.002731 | 0.001433 | 0.0003139 |
|  | R<sup>2</sup> | 0.9738 | 0.9667 | 0.9927 |
|  | Adjusted R<sup>2</sup> | 0.9679 | 0.9594 | 0.9886 |
|  | RMSE | 0.01742 | 0.01262 | 0.006696 |

**Figure 4.**
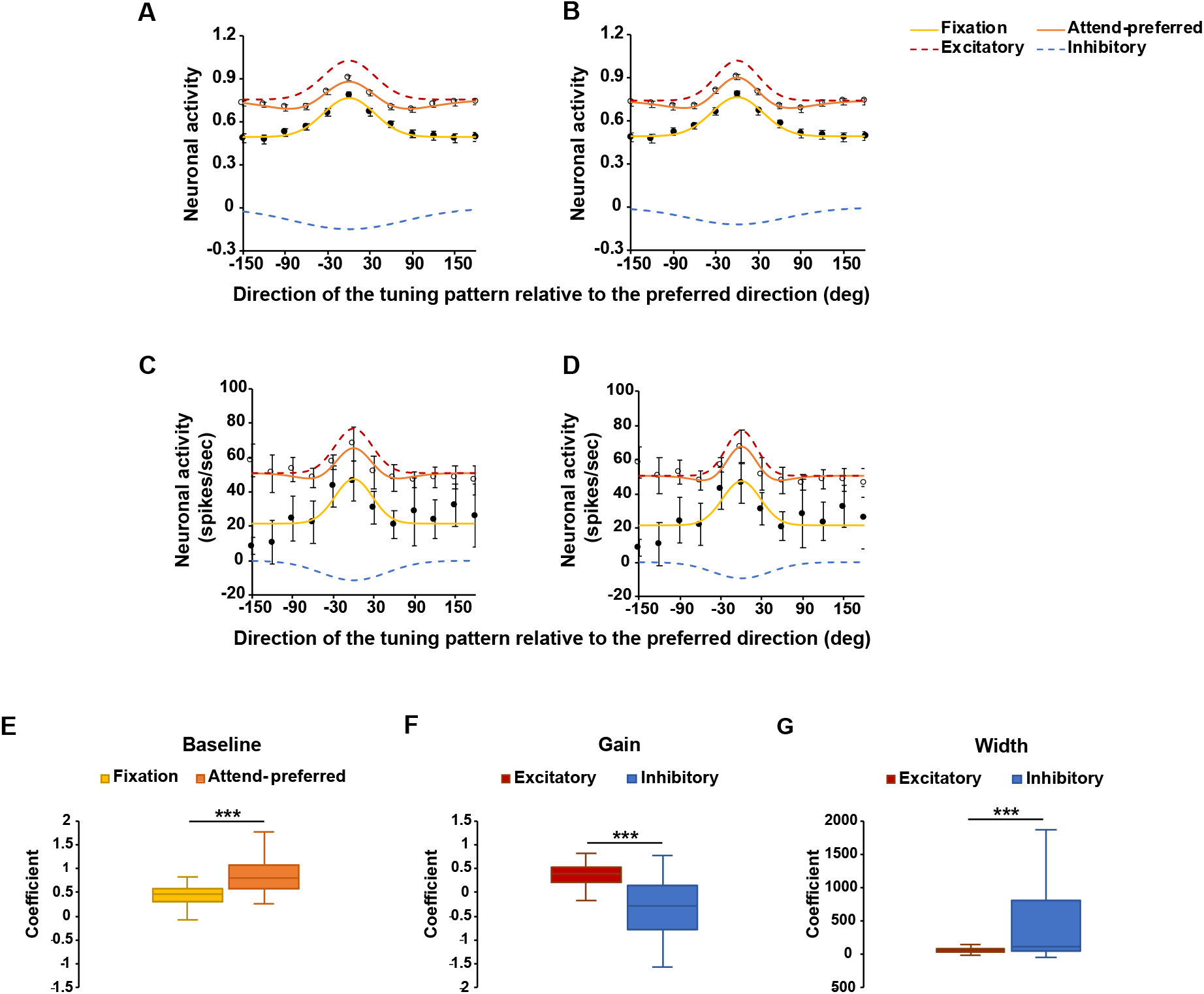
(A) Model fits to the averaged normalized neuronal responses (outlined circles: attend-preferred, filled circles: fixation). For the sum of two Gaussians model in the attend-preferred condition, coefficients of the excitatory Gaussian (gain and width) were the same as those of the single Gaussian model in the fixation condition. (B) All coefficients of the sum of two Gaussians model were free to vary. (C-D) Examples of the model fits to individual neuronal responses. In (C), the coefficients of the excitatory Gaussian were constrained while they were unconstrained in (D). (E-G) Median coefficients of the model fits to the normalized individual neuronal data. (E) Baseline coefficients when the single Gaussian model fit the data in the fixation condition and when the sum of two Gaussians model fit the data in the attend-preferred condition. (F) Gain and (G) width coefficients of the excitatory and inhibitory Gaussians when the sum of two Gaussians model fit the data in the attend-preferred condition. Note that the gain coefficients of the excitatory Gaussian were the same as those of the single Gaussian fits in the fixation condition. Error bars in Fig 4A-B, and 4E-G indicate SEM and those in Fig 4C-D indicate SD. *** *p* <. 001

Second, we fit the average normalized neuronal response in the attend-preferred condition with a sum of two Gaussians model. We fixed the parameters of one Gaussian to the same parameters obtained from the fit of the fixation condition. This would approximate the excitatory field provided by the tuned inputs into the cell. For the second Gaussian, the coefficients were not constrained. We assumed that this second Gaussian would have negative gain which represents inhibitory-tuned inputs into the neuron recruited (or magnified) by FBA. The equation is:

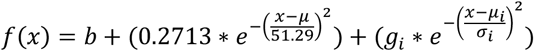

The parameters *b* (baseline), *g*_*i*_ (gain), and *σ*_*i*_ (width) were free to vary. The baseline was not fixed since it may capture overall changes in response due to spatial attention or arousal, i.e., in the fixation condition attention was directed to the fixation point while in the attend-preferred condition attention was directed to the RF and attending to the RF produces a response increase for all directions ^8^. The other parameters, *g*_*i*_ and *σ*_*i*_ represent the gain and width of the inhibitory inputs into a neuron. Negative values of *g*_*i*_ would reflect increase in the gain of inhibitory inputs/field by attention.

In the attend-preferred condition, the baseline of neuronal response increased (red dashed line) relative to the baseline in the fixation condition (yellow line) (Figure 4A and Table 1). This may reflect the increase in responses due to directing attention into the RF. Importantly, as anticipated, the second Gaussian which represents the inhibitory field (blue dashed line) showed a negative gain (*g*_*i*_ < 0) and a broader (*σ*_*i*_ > *σ*) width than the first Gaussian representing the excitatory field (51.29 vs. 110.8, Table 1). Notice the excitatory field was estimated from the Gaussian fit in the fixation condition, where we assume the contribution of the inhibitory field was small relative to that of the excitatory field.

One might argue that our estimation of the excitatory field may have not been accurate and that both, the shapes of the excitatory and inhibitory fields will change in the attend-preferred condition (e.g., a narrowing of the excitatory field is plausible). In order to test this, we repeated the fitting procedure but letting the coefficients of both Gaussian functions freely vary (Figure 4B). Interestingly, the improvement in the goodness-of-fit of the model and changes in the coefficients of both Gaussians were small (Table 1). This indicates that our estimation of the excitatory field is reasonable, and that changes induced by FBA on the inhibitory field alone can account for the observed changes in tuning profile.

In order to further corroborate this result, we repeated a similar procedure at the level of single neurons. We first fit the single Gaussian model to the normalized individual neuronal responses in the fixation condition. The coefficients of each individual fit were used to model the same neuron’s excitatory field in the attend-preferred condition as we did in the population level analysis. The coefficients for the other Gaussian, which models the inhibitory field, were not constrained. Neurons were excluded from the analyses if the fitting procedure was not successful due to missing data points or severe variability in the data (i.e., fit does not converge). Consequently, 69 neurons were included in the analysis. The median baseline, gain, and width coefficients of the single Gaussian model fit in the fixation condition were 0.4656, 0.3996, and 54.83, respectively. In the attend-preferred condition, when the excitatory Gaussian coefficients (gain and width) were constrained (same as those of the single Gaussian fits in the fixation condition), the median baseline was 0.808 and the median gain and width of the inhibitory Gaussian were −0.2889 and 113.97, respectively. Wilcoxon signed-rank tests showed that the baseline was significantly elevated in the attend-preferred condition (*Z* = 4.1105, *p* < .001, Figure 4E). The inhibitory fields had significantly lower gains (*Z* = −3.9371, *p* < .001, Figure 4F) and the broader widths (*Z* = 5.0612, *p* < .001, Figure 4G) than the excitatory fields. This shows that model fits to the individual neuron data show the same results as in the population level analysis.

### Behavioral effects of feature-based surround suppression

We investigated the effect of feature-based surround suppression on behavior using motion repulsion ^23–25^. Motion repulsion was measured when participants divided their attention to two superimposed motion surfaces (divided attention condition) or focused on one of them (focused attention condition, Figure 5A). The superimposed motion surfaces were separated by different colors and their directional offsets systematically varied (10 to 50 deg). The speed of motion for the surfaces could be 3 deg/sec or 6 deg/sec.

**Figure 5.**
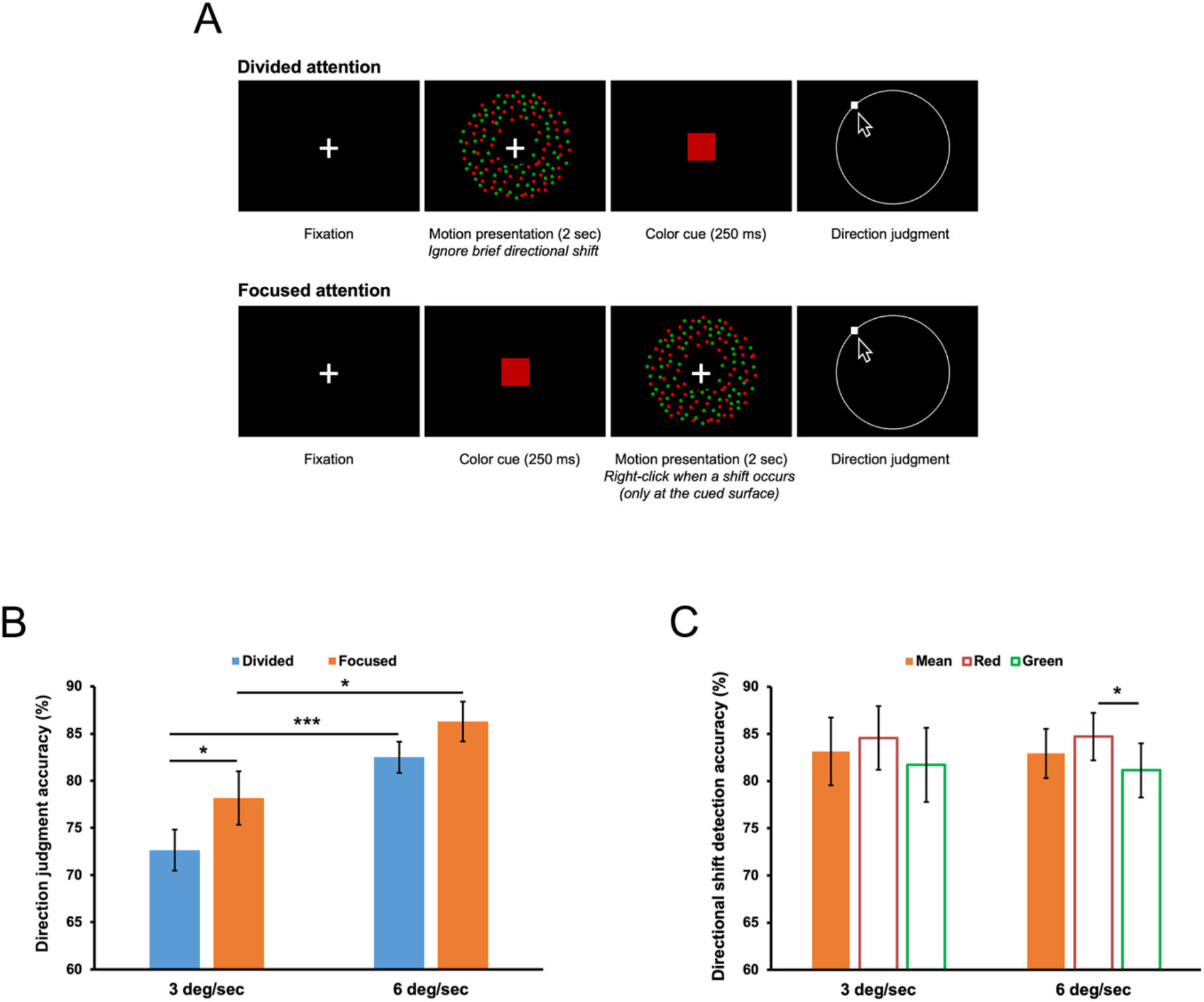
(A) Experimental conditions and procedure of human behavioral experiment. In the divided attention condition, participants attended to both motion surfaces equally, ignoring any directional shift. A color cue was presented after RDPs disappeared and then, participants reported a direction of the cued motion surface by clicking along a white circular outline. In the focused attention condition, a color cue indicated which motion surface participants had to attend. Participants were asked to detect a directional shift on the cued surface if it occurred, and then reported motion directions of this surface as in the divided attention condition. (B) Direction judgment was more accurate in the focused attention condition, and when motion speed was faster. (C) Detection of directional shifts (only in the focused attention condition) was not affected by motion speed. However, direction discrimination was more accurate when the attended motion surface was red. Error bars indicate SEM. * p < .05, *** p <. 001

First, we analyzed how FBA and motion speed influenced participants’ direction judgment accuracy – i.e., whether reported directions fell within a valid response range (Figure 5B, see Data analysis in Methods). Direction judgment accuracy in the focused attention condition was measured only if the attentional task (detecting a brief directional shift) was successfully performed. Bonferroni correction was used for multiple comparisons. A repeated-measures ANOVA demonstrated significant main effects of the attentional conditions (*F*(1, 13) = 6.683, *p* = .023) and motion speed (*F*(1, 13) = 22.443, *p* < .001) on direction judgment accuracy. Post-hoc pairwise comparisons showed that direction judgment was more accurate in the focused attention condition than in the divided attention condition (M_diff_ = 4.59%, SE = 1.78%) for both motion speeds (3 deg/sec: *p* = .041, 6 deg/sec: *p* = .065 (trend)) because participants tracked only one motion direction in the focused attention condition. Direction judgment accuracy was also higher when motion speed was faster (M_diff_ = 9.03%, SE = 1.91%) in both attentional conditions (divided: *p* < .001, focused: *p* = .011). Color of the target motion surface did not influence direction judgment (*F*(1, 13) = .051, *p* = .825). All interactions between the variables were not significant.

Figure 5C shows the mean accuracy of directional shift detection in the focused attention condition. The mean accuracy was 83.14% (SD 13.45%) and 82.93% (SD 9.79%) when motion speed was 3 deg/sec and 6 deg/sec, respectively. They did not statistically differ (*p* = .915), indicating that difficulty of the attentional task was well controlled across different speed conditions. When it was broken down by the color of the attended motion surface (target), the main effect of the target surface color on directional shift detection was significant (*F*(1, 13) = 13.995, *p* = .002). Detecting directional shifts was better when the target motion surface was red than when it was green (M_diff_ = 3.21%, SE = .86%) for both motion speeds (3 deg/sec: *p* = .062 (trend), 6 deg/sec: *p* = .015). The performance advantage in the red motion surfaces likely represents strong attentional guidance of red ^29,30^. Interaction between motion speed and the color of the target motion surface was not significant (*F*(1, 13) = .12, *p* = .734).

We analyzed motion repulsion of the trials in which participants correctly discriminated motion direction. When motion speed was 3 deg/sec (Figure 6A), the main effect of the attentional condition on motion repulsion was not significant (*F*(1, 13) = 2.224, *p* = .16), whereas motion repulsion significantly varied depending on the directional difference between the two motion surfaces (*F*(4, 52) = 2.715, *p* = .04). Importantly, motion repulsion was influenced by the interaction between the attentional condition and directional difference (*F*(4, 52) = 3.958, *p* = .007). Motion repulsion was significantly reduced in the focused attention condition when the directional difference was 30 deg (M_diff_ = 4.40 deg, SE = 1.41 deg, *p* = .008) and this effect was marginal when directional difference was 40 deg (M_diff_ = 2.81 deg, SE = 1.34 deg, *p* = .056). Reduction of motion repulsion at around 30~40 deg difference suggests that feature-based surround suppression played a role by inhibiting an unattended motion direction that was near the attended motion direction.

**Figure 6.**
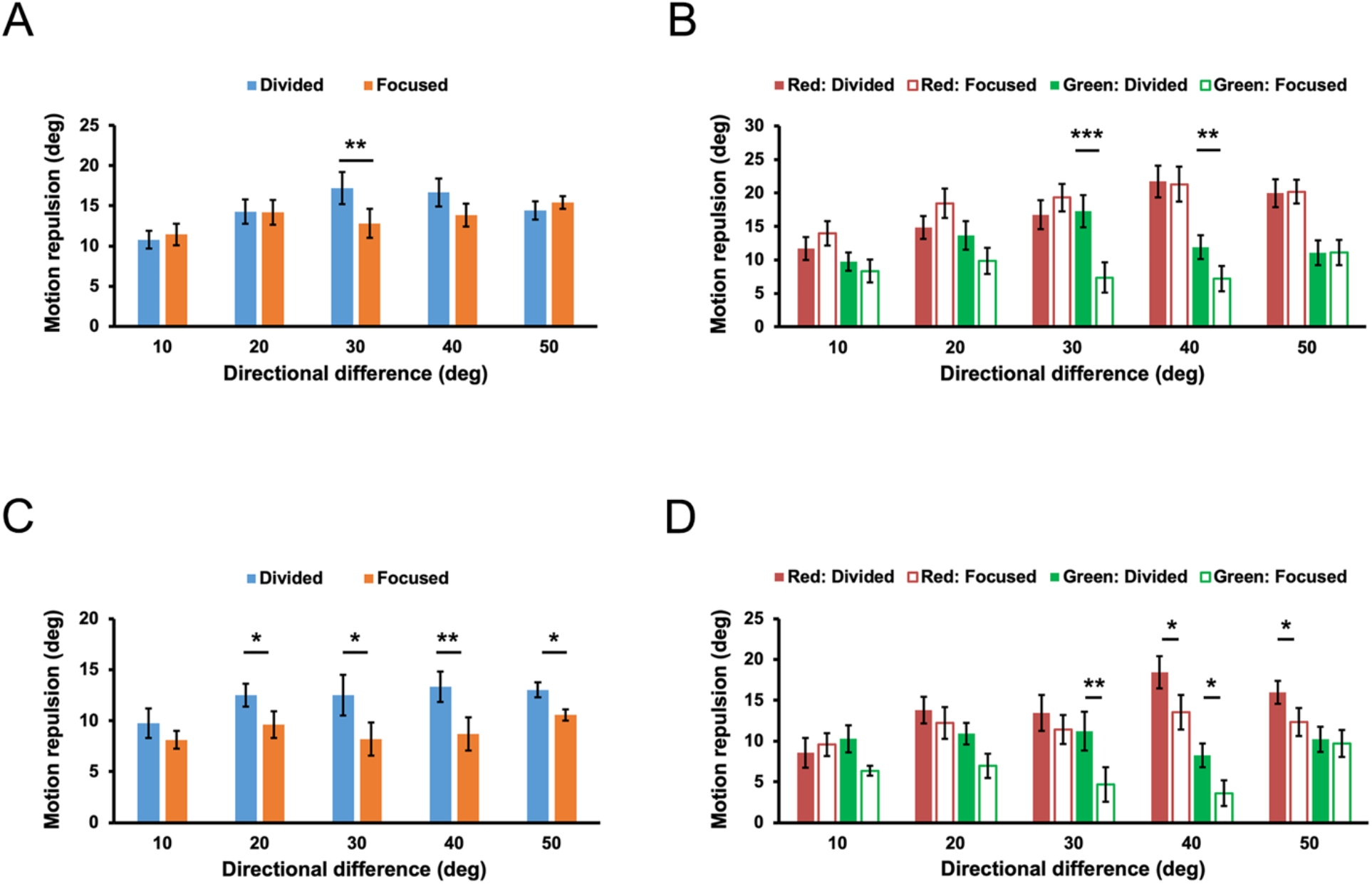
When motion speed was 3 deg/sec, (A) the amount of motion repulsion between the two attentional conditions significantly differed when directional difference was 30 deg. The same effect was marginal when directional difference was 40 deg (*p* = .056). (B) Attentional modulation on motion repulsion was evident only when the color of the target motion surface was green. When motion speed was 6 deg/sec, (C) Motion repulsion was reduced in the focused attention condition and it was true across all directional differences, except at 10 deg difference. (D) Unlike when motion speed was slower, attentional modulation on motion repulsion was observed regardless of the color of the target motion surface. Error bars indicate SEM. * *p* < .05, ** *p* < .01, *** *p* < .001

We further broke down the data by the color of the target motion surface to see how this factor is associated with motion repulsion. The amount of motion repulsion was different depending on the color of the target motion surface (*F*(1, 13) = 28.589, *p* < .001; Figure 6B). Motion repulsion was smaller when the target motion surface was green than when it was red (M_diff_ = 7.07 deg, SE = 1.32 deg). Although the attentional condition did not affect motion repulsion (*F*(1, 13) = 2.07, *p* = .174), there was a significant interaction between the target motion surface color and the attentional condition (*F*(1, 13) = 9.199, *p* = .01). We conducted an ANOVA separately for each target surface color to investigate this interaction more specifically. When the target motion surface was red, motion repulsion was not affected by the attentional conditions (*F*(1, 13) = 1.188, *p* = .296) but by the directional difference between the two motion surfaces (*F*(4, 52) = 7.522, *p* < .001). The interaction between these two variables was not significant (*F*(4, 52) = .8, *p* = .531). Hence, there was no evidence for reduction of motion repulsion by FBA and feature-based surround suppression. On the other hand, when the target motion surface was green, the main effect of the attentional conditions was significant (*F*(1, 13) = 22, 631, *p* < .001), indicating that FBA reduced motion repulsion (M_diff_ = 3.96 deg, SE = .83 deg) as previously reported ^26^. Motion repulsion was not modulated by directional difference (*F*(4, 52) = .802, *p* = .529) but the interaction between the attentional conditions and directional difference was significant (*F*(4, 52) = 6.699, *p* < .001). Motion repulsion significantly decreased in the focused attention condition when directional difference was around at 20~40 deg (20 deg: *p* = .051 (marginal), 30 deg: *p* < .001, and 40 deg: *p* = .005). This indicates that feature-based surround suppression in the 3 deg/sec condition was mainly derived from the trials where the target motion surface was green.

Change in motion speed from 3 deg/sec to 6 deg/sec affected motion repulsion (*F*(1, 13) = 54.343, *p* < .001) – faster motion speed (6 deg/sec) reduced motion repulsion (M_diff_ = 3.47 deg, SE = .47 deg) as previously reported ^31–33^. When motion speed was 6 deg/sec (Figure 6C), the attentional condition significantly modulated motion repulsion (*F*(1, 13) = 19.557, *p* = .001), demonstrating that motion repulsion was generally smaller in the focused attention condition (M_diff_ = 3.18 deg, SE = .72 deg). Directional difference (*F*(4, 52) = 1.652, *p* = .175) and the interaction between the attentional condition and directional difference (*F*(4, 52) = 1.049, *p* = .391) did not significantly affect motion repulsion. Motion repulsion was significantly smaller in the focused attention condition at all directional differences, except at 10 deg difference, and the difference in motion repulsion (divided - focused) did not statistically vary across directional differences. This result made it difficult to specify the effect of feature-based surround suppression.

Motion repulsion was modulated by the color of the target surface (*F*(1, 13) = 14.491, *p* = .002), showing smaller motion repulsion in green surfaces (M_diff_ = 3.07 deg, SE = .76 deg; Figure 6D). In addition, the attentional condition significantly affected motion repulsion (*F*(1, 13) = 16.507, *p* = .001). Motion repulsion was smaller in the focused attention condition than in the divided attention condition (M_diff_ = 3.07 deg, SE = .76 deg). The interaction between the target surface color and the attentional condition was not significant (*F*(1, 13) = .865, *p* = .369). Unlike when motion speed was 3 deg/sec, the attentional condition modulated motion repulsion even when the color of the target motion surface was red (*F*(1, 13) = 5.177, *p* = .04). Post-hoc pairwise comparisons showed that motion repulsion was reduced in the focused attention condition (M_diff_ = 2.22 deg, SE = .97 deg) and it was significant when the directions of RDKs differed by 40~50 deg (40 deg: *p* = .036, 50 deg: *p* = .02). When green was the color of the target motion surface, FBA also significantly reduced motion repulsion (*F*(1, 13) = 8.148, *p* = .014; divided vs. focused: M_diff_ = 3.93 deg, SE = 1.38 deg). The reduction was at the trend level at 10~20 deg difference (10 deg: *p* = .053, 20 deg: *p* = .085) and became significant at 30~40 deg difference (30 deg: *p* = .008, 40 deg: *p* = .021). Motion repulsion patterns for green motion surfaces were consistent across different motion speeds, whereas FBA played a role only if motion speed was faster when the target surface was red. Therefore, the almost universal decrement in motion repulsion in the 6 deg/sec condition might have resulted from the interaction between the color of the target motion surface and motion speed.

## Discussion

The present study demonstrated feature-based surround suppression in the motion direction domain at both neurophysiological and behavioral levels. The neurophysiological findings show that attention to a neuron’s preferred motion direction modulates its direction tuning curve, inducing a suppressive surround around the attended direction. In addition, our behavioral study demonstrates feature-based surround suppression on motion repulsion in humans.

### Effects of FBA on motion direction tuning curves

FBA allows enhancing the processing of attended features while suppressing the processing of unattended ones ^5^. These effects of FBA have been captured by the feature-similarity gain model, which proposes that modulation of a neuron’s response is a monotonic function of the differences between the neuron’s preferred feature and the attended feature ^4,8^. FBA produces a maximal response enhancement for the attended feature and a progressive decrease of responses that becomes suppression relative to a neutral condition as the attended feature deviates from the preferred feature of the neurons. Our results in this study do not match those reported by previous electrophysiological studies of FBA in macaques ^4,8^. We found that the decay in the intensity of the FBA response enhancement was not monotonic but reached its minimum in the vicinity of the attended feature.

One possible explanation for the difference between our results and those of a previous study is that in the aforementioned study^4^ there was a large separation between the target and the distractor (i.e., located in opposite hemifields), therefore the FBA response modulation was small and may not have been sufficient to estimate the precise shape of the feature-similarity modulation. Location-based surround suppression predicted by ST was formulated for stimulus conditions in which distractors are nearby the attended stimulus and attention filters out their contribution to the neuron’s response. In the present study the attended stimulus and distractor shared the same RF, they were nearby, resembling the circumstances under which ST was formulated. FBA modulation was stronger because the proximity of the RDPs may have triggered stronger inhibitory circuit dynamics. Indeed, it has been reported that attentional modulation is stronger when targets and distractors are positioned inside the RF relative to when they are positioned one inside and the other outside the RF ^8,34^. Moreover, distractor interference is stronger when both stimuli are within the same relative to different hemifields ^35^. One additional detail is that in our study we maintained the preferred pattern in the RF, which may have strongly driven the responses of the recorded neurons improving signal to noise ratio ^36^ in a way that the modulation was detectable with a reasonable sample of recorded neurons.

### Modeling the response modulation

Models of MT neurons have proposed that feature tuning is mainly a function of excitatory-tuned inputs into a neuron that are modulated by inhibitory inputs from pools of neurons within the same area. This idea is captured by normalization models ^37^. However, in normalization models, the strength of inhibitory inputs into a neuron does not depend on the stimulus feature. Normalization models have been proposed to account for the shape of the modulation of neuronal responses by attention ^38^. Attention could act via modulating inputs into neurons, because the same inputs activate the normalization pool in different manners depending on the size of the attentional focus, a variety of effects could be achieved at the level of single cell responses. Others have further proposed tuned normalization as a mechanism to explain the variety of attentional modulation across different neurons ^39,40^. In the latter studies, the normalization pool is tuned by the attentional modulation, or an existing tuning is modulated by attention.

One possibility that explains the effects we report here is that a tuned normalization pool is modulated by feedback signals from high-order areas into MT/MST. However, for a gain feedback modulation to explain the feature-based surround suppression, the tuning of the feedback modulation, or of the inhibitory neuronal pools they activate directly or indirectly in MT/MST needs to be wider than the tuning of feedforward excitatory inputs. It is possible the feedback does not directly drive inhibitory neurons but excitatory cells that in turn activate the inhibitory pool. It is difficult with the available data to provide a detailed circuit layout. But we can elaborate on a hypothesis as follows: 1) the tuning/width of the inhibitory inputs into a neuron are wider than those of excitatory inputs, and b) attention modulates the gain of inhibitory inputs more strongly than excitatory inputs. The appeal of this proposal relies on how gain changes differentially applied to MT/MST excitatory and inhibitory fields could produce a non-monotonic modulation. We should note that feedforward inputs into a neuron could increase their strength with attention as long as the effect is smaller than the one of feedback inputs. Indeed, this may be the case according to studies isolating the modulation of responses in area V1 ^34^. If feedback gain signals were to originate downstream from the recorded area, and given that the attentional modulation of responses grows along the hierarchy of visual processing ^41,42^, it is reasonable to assume that within a given area changes in the strength of feedback signals would be greater than changes in the strength of feedforward inputs.

Some studies using calcium imaging and genetically labelled cell types in mice have reported that the tuning of inhibitory neurons for stimulus features is wider than the tuning of excitatory cells ^43^. In monkey MSTd (one of areas we recorded neuronal responses from in this study) narrow-spiking putative interneurons are more broadly direction tuned than broad-spiking putative pyramidal cells ^44^. It is reasonable to assume that narrow-spiking cells are in their majority inhibitory interneurons while broad-spiking cells are in their majority pyramidal cells ^45^. Moreover, given that the PV cell is the most abundant interneuron type in MT/MST ^45^, and it is involved in gain control of pyramidal cells ^46^, one could elaborate on the hypothesis that PV cells recruited by top-down inputs driven by attentional templates of attended features ^47^ provide the wide strong inhibitory drive that produces the non-linear change in the overall response curve. Interestingly, previous studies of attention in area MT reported that the widths of tuning curves with attention tend to be wider (although not statistically different) than those in unattended conditions ^8^. This may suggest that the wider tuning of inhibitory fields may be at least influenced by feedback signals. Whether changes in tuning of narrow spiking cells in MT/MST could be induced by attention or it is intrinsic to the area circuitry needs to be further investigated.

Finally, ST predicts the effects described here as resulting from top down signals. Interestingly, studies using moving RDPs have shown that the latency of the attentional effects on the responses of direction-selective neurons is shorter in LPFC than in MT ^42^, suggesting that top-down FBA signals originate in executive areas and feedback into visual cortex to modulate processing ^47,48^. Our study reveals additional pieces of information that can be incorporated into ST to generate more specific predictions at microcircuit level.

### Psychophysics

We observed that attention to one of the two superimposed RDKs reduced motion repulsion and this reduction was greatest when the two RDKs moved in similar directions, suggesting feature-based surround suppression. This suppressive effect varied depending on bottom-up factors such as motion speed and the color of the target motion surface which indicates an interplay between top-down FBA and bottom-up factors. The present results appear to be inconsistent with the previous report that the color of motion surfaces does not influence motion repulsion ^49^. However, in their study, motion repulsion was not measured separately for the motion surface colors, hence, the effect of the surface color might have not been addressed. Greater motion repulsion in red surfaces suggests that these surfaces are more strongly affected by inhibition. Such a result is consistent with a recent study that showed red facilitated response inhibition compared to green ^50^. It is also possible that different color wavelengths may influence motion repulsion. When the speed of the motion was faster, producing more signal strength, it is likely that the balance in mutual inhibition between the two-colored surfaces was more even, which then allowed for feature-based attentional modulation and surround suppression to be seen on the red surface.

One puzzling finding in the present research was that the feature distance where surround suppression occurred was different across experiments. Feature-based surround suppression in our behavioral experiment in humans was produced when the directional difference between two motions was smaller (30~40 deg) than in the neurophysiological experiment in monkeys (60~90 deg). The previous behavioral studies reported broader feature-based suppressive surrounds in the motion direction dimension than those in the other feature dimensions (maximum suppression around at around 90 deg difference, ^20,21^) and this range is similar to what we found in the neurophysiological experiment, suggesting the broad direction tuning curve of MT neurons ^51–53^. There are several possible explanations for the discrepancy between our two experiments. It may be that responses of MT single neurons tuned for motion direction changed into a different population response profile in areas toward which MT projects (e.g., LIP or the lateral prefrontal cortex). In favor of this hypothesis, it has been reported that during a task that requires categorization of motion directions, LIP and prefrontal neurons change their tuning while MT neurons do not ^54^. It may also be that a narrower surround suppression profile in our behavioral experiment could be due to the nature of motion repulsion. Motion repulsion is typically attenuated as the two superimposed motions move in more dissimilar directions, thus, one would not find feature-based surround suppression when the directional difference becomes greater. In addition, human participants were asked to report motion directions which requires higher precision of perception than the monkeys’ task where they only had to detect directional changes while maintaining attention on an eccentric motion pattern. Finally, we used colored, superimposed RDPs in the behavioral experiment, while we used RDPs of the same color and spatially separated in the neurophysiological experiment. These differences in experimental designs and possible biases idiosyncratic to each species (humans vs. monkeys) may influence the profile of feature-based surround suppression.

Compared to the previous studies ^20,21^, the current study provided direct and stronger evidence of surround suppression in direction discrimination. In the prior studies, participants were required to detect a coherent motion signal, rather than reporting the direction of motion, which was the domain for feature-based surround suppression being investigated. In Wang et al.^21^, the experimental design also separated all stimuli temporally, potentially confounding visual working memory and FBA. Hence, their results would be inferential on effects of motion direction processing. In contrast, the current study used real motion to initiate FBA and required participants to discriminate the direction of motion. In addition, the stimuli in this study were superimposed spatially and temporally. Together, the current behavioral paradigm allowed for directly testing feature-based surround suppression in real-time active circuitry. As motion repulsion is thought to occur due to mutual inhibition between competing pools of direction-selective neurons^55^, inhibiting the neurons representing directions close to the attended direction in feature space through feature-based surround suppression would then reduce inhibition on the neurons representing the attended direction. This shifts the population tuning back towards the veridical direction from the repulsed one, consistent with our results and ST. Therefore, our experimental paradigm more directly demonstrates feature-based surround suppression of ST relative to the previous ones.

## Conclusion

Our results demonstrate that feature-based attention is a flexible mechanism capable of modulating neuronal responses in the visual cortex. Moreover, it shows how non-linearities in the effects of feature-based attention observed during neurophysiological and psychophysical studies can emerge from the interaction between independent gain control mechanisms on excitatory and inhibitory elements of a microcircuit that contribute to a neuron’s response. Ultimately, our results show how feature-based attention uses the intrinsic architecture of cortical microcircuits to enhance the processing of targets at the expense of distractors.

## Methods

### Neurophysiology in macaque monkeys

Research with non-human primates represents a small but indispensable component of neuroscience research. The researchers in this study are aware and are committed to the responsibility they have in ensuring the best possible science with the least possible harm to the animals ^56^.

#### Apparatus and stimuli

We recorded the responses of direction-selective neurons in areas MT and MST (n = 107) of two male adult macaque monkeys in different task conditions. After initial training, a head post, a scleral search coil ^57^ to monitor eye position ^58^, and a recording chamber were implanted in each animal. A custom computer program running on an Apple Macintosh PowerPC controlled the stimulus presentations, monitored eye position and behavioral responses during the experiments, and recorded the behavioral and neuronal data. The experiments reported in this study were conducted according to local and national rules and regulations and were approved by the Regierungspraesidium Tuebingen (Germany).

RDPs consisted of small bright dots with a density of 5 dots/dva^2^ within a stationary circular virtual aperture on a dark computer monitor. The luminance of the dots was 55 cd/m^2^ and the viewing distance was 57 cm. The diameter of the aperture varied from about 1 deg to 12 deg depending on the size of each neuron’s RF so that the stimulus did not exceed the boundaries of the classical RF. Movement of the dots was created by displacement of each dot by the appropriate amount at the monitor refresh rate of 75 Hz. We measured the speed and direction tuning of the neuron by online display of the tuning curves and chose the speed at which the neuron produced the strongest response ^8^. In every trial we presented two RDPs of equal size inside the neurons RF. One RDP always moved in the neuron’s preferred direction, which was estimated in a separate block of trials by online display of the responses to a single RDP inside the RF moving in different directions while the animal was fixating a dot at the screen center (see ^8^). The other pattern could move in one of twelve different directions spaced every 30 deg.

#### Recordings

Extracellular recordings from the left hemisphere were conducted using tungsten microelectrodes (impedance 0.5–2 mΩ, Microprobe and FHC). Electrodes were lowered through a recording chamber implanted on top of the parietal bone until reaching the approximate location of MT/MST. Single units were isolated with a window discriminator, and eccentricity, direction-selectivity, and position of the electrode within the recorded area were determined. We recorded only from those neurons showing clear direction-selectivity during initial mapping (see ^8^ for more detail). Throughout this process, we recorded from 107 MT/MST neurons (75 MT and 32 MST neurons).

#### Task

The animals were trained to selectively attend to a cued stimulus, while directing gaze to a fixation point (Figure 1A). One of the RDPs always moved in the neuron’s preferred direction (preferred pattern) and the other could move in one of 12 different directions from trial to trial (tuning pattern, in steps of 30 deg). There were three different experimental conditions depending on which feature on the display was attended (Figure 1B). The animals were cued to attend to 1) the preferred pattern (attend-preferred condition), or 2) tuning pattern (attend-tuning condition). As an attentional cue, the target appeared for a short time period and then disappeared to re-appear together with the distractor. The task for the animals was to covertly attend to the target pattern and release a lever within a 450 ms response window after the dots changed direction. The animals had to ignore changes in the direction of the unattended pattern, which happened in 50% of the trials. 3) In a third condition (fixation condition), the animals attended to the fixation point and released a lever within a 450 ms after the fixation point changed color while ignoring both RDPs in the periphery.

#### Data analysis

Neurons were included in the analysis if the number of data points in which they were recorded from was more than 6. As a result, 78 MT/MST neurons were analyzed (53 MT and 25 MST neurons). We computed average firing rates during the interval from 200 to 1200 milliseconds after the onset of the two patterns, as a function of the tuning pattern’s direction relative to the direction of the preferred pattern. The responses of each neuron were normalized to the response when both RDPs moved in the neuron’s preferred direction in the attend-preferred condition and then averaged across neurons. The responses of both MT and MST neurons were pooled since the direction selectivity and tuning curve profiles were very similar between the two areas. Repeated-measures ANOVA and paired samples t-tests on the average normalized neuronal response were conducted to examine how neuronal responses varied depending on experimental conditions.

We fitted the single Gaussian and sum of two Gaussians models to neuronal responses, using the MATLAB curve fitting toolbox (Mathwork Inc., USA). The sum of two Gaussians model we used is equivalent to the difference-of-Gaussians (DoG) model as the first Gaussian has a positive gain and the second Gaussian has a negative gain. We used the Akaike Information Criterion (AIC, ^59^) which penalizes model complexity (lower AIC value indicates better fit) to assess the relative quality of each model fit.

### Human behavioral experiments

#### Participants

Fourteen naïve participants (5 men, 9 women), between the ages of 20 and 30 years completed the experiment. They had normal or corrected-to-normal visual acuity, and normal color vision. Written informed consent was obtained from all participants and they were paid for their participation ($30 CAD per subject). The research was approved by York University’s Human Participants Ethics Review Committee.

#### Apparatus and stimuli

Experiments were conducted in a dark room. Participants sat 57 cm from a CRT monitor (21” View Sonic G225f, 1280×1024, 85 Hz) and their heads were stabilized on a head and chin rest (Headspot, UHCOtech, Houston, TX). Participants wore an infrared eye tracker (Eyelink II, SR Research, 500 Hz, Mississauga, ON, Canada) monitoring the left eye position. Random dot patterns (RDPs) were created through MATLAB (MathWorks, Natick, MA) and the Psychophysics Toolbox ^60,61^. Experimental control was maintained by Presentation (Neurobehavioral Systems, Berkeley, CA).

An annular RDP consisted of two superimposed motion surfaces (RDP size = 15 dva (degree in visual angle) in diameter, inner aperture size = 6 dva in diameter, dot size = 0.15 dva, 75 dots per surface). The directions of the motion surfaces changed every trial and the dots in each motion surface moved in the same direction (100% coherent). Directional difference between the two surfaces systematically varied by 10~50 deg (10 deg step). Motion speed was either 3 deg/sec or 6 deg/sec and both motion surfaces moved in the same speed. Dots in one motion surface were red (luminance: 24.67 cd/m^2^) and those on the other surface were green (24.64 cd/m^2^) to make participants easily segregate them, without affecting direction repulsion ^49^.

#### Task

We tested participants under two experimental conditions: divided and focused attention. In the divided attention condition, typical motion repulsion was measured. The RDPs were presented for 2 sec once participants fixated a white cross centered on a screen for 200 ms. Participants had to maintain the fixation until the RDPs disappeared, otherwise an error message was presented, and the trial was randomly interleaved in the remaining trials. Participants were asked to view both motion surfaces equally to estimate their directions. During motion presentation, a brief directional shift (100 ms) on either motion surface could randomly occur in 80% of trials, and then it went back to the original direction. The amount of shift was randomly selected from the range between 30 and 40 deg when motion speed was 3 deg/sec, and between 20 and 30 deg when motion speed was 6 deg/sec to equalize the perceptual strength of directional shifts across different motion speed conditions. A shift could occur from 650 to 1100 ms after RDPs onset. Participants were asked to ignore this directional shift while they viewed the RDPs. After motion presentation, a color cue (either red or green) appeared for 250 ms to indicate which motion surface was the target in that trial. Each color cue was presented equally throughout the experiment. Participants reported the motion direction of the target surface by clicking along a white circular outline.

In the focused attention condition, after maintaining central fixation for 200 ms, a color cue appeared before the RDPs were presented to indicate which motion surface should be attended. Participants were required to attend only to the cued motion surface (target) while ignoring the other surface. To make sure whether they selectively attended to the target surface, they clicked the right mouse button within a 1 sec after the onset of a brief directional shift. The directional shift could occur only on the target surface. If there was no shift, participants did not respond and waited until they viewed the white circular outline. If participants missed the shift, responded too late, or made a false alarm, an error message was presented, and the trial was discarded. They reported the motion direction of the target surface only when the attention task was successful performed.

The attentional conditions were blocked, and participants performed both conditions twice in a random order. At the beginning of each attentional condition, participants were given 10 practice trials whose data were not used and then, they performed 100 trials as the main experiment (200 trials for each attentional condition, in total). There was a mandatory “break time” after every 25 trials. Participants could have extra “break time” if they wanted.

#### Data analysis

We first sorted participants’ direction judgment responses to reduce variability in data ^49^. A correct direction judgment should fall within a range that extended from halfway between the two motion directions to 45 deg away from the motion direction of the target surface. Then, motion repulsion, defined by the difference between the reported (perceived) and the actual motion direction, was calculated. Since participants performed an additional attention task in the focused attention condition, only trials in which both directional shift detection and motion direction judgment were successful were included in the analysis. Motion repulsions in the two attentional conditions were compared to quantify the attentional modulation.

## Data availability

Data and codes of the current study are available at https://osf.io/5qn74/

## Acknowledgements

The authors thank Daniel Voloshin and Amir Yazdanparast for helping human data collection. S.-A.Y. and J.K.T. are funded by Air Force Office of Scientific Research (FA9550-18-1-0054), the Canada Research Chairs Program (950-231659), and the Natural Sciences and Engineering Research Council of Canada (RGPIN-2016-05352). J.M.T. is funded by the Canadian Institutes for Health Research Operating Grant. S.T. is funded by the Deutsche Forschungsgemeinschaft (DFG, German Research Foundation) for the Collaborative Research Center 889 “Cellular Mechanisms of Sensory Processing” (Project C04) and the Research Unit 1847 “Physiology of Distributed Computing Underlying Higher Brain Functions in Non-Human Primates” (Project A1). M.F. is funded by the Natural Sciences and Engineering Research Council of Canada (RGPIN-2016-05296).

## Author contributions

J.M.T. and S.T. designed and planned the neurophysiological experiment.

J.M.T. performed the neurophysiological experiment.

S.-A.Y., J.K.T. and M.F. designed and planned the psychophysical experiment.

S.-A.Y. performed the psychophysical experiment.

S.-A.Y and J.M.T. analyzed the data.

All the authors contributed to the interpretation and discussion of the results, and wrote the manuscript.

